# A scalable method for automatically measuring pharyngeal pumping in *C. elegans*

**DOI:** 10.1101/051243

**Authors:** Monika Scholz, Dylan J. Lynch, Kyung Suk Lee, Erel Levine, David Biron

**Affiliations:** James Franck Institute, the University of Chicago, Chicago, IL 60637; Institute for Biophysical Dynamics, the University of Chicago, Chicago, IL 60637; Department of Chemical Engineering, University of Illinois at Chicago, Chicago, Illinois 60607; Department of Physics and Center for Systems Biology, Harvard University, Cambridge, MA 02138; Department of Physics, the University of Chicago, Chicago, IL 60637^*^

## Abstract

We describe a scalable automated method for measuring the pharyngeal pumping of *Caenorhabditis elegans* in controlled environments. Our approach enables unbiased measurements for prolonged periods, a high throughput, and measurements in controlled yet dynamically changing feeding environments. The automated analysis compares well with scoring pumping by visual inspection, a common practice in the field. In addition, we observed overall low rates of pharyngeal pumping and long correlation times when food availability was oscillated.

## I. INTRODUCTION

The nematode *Caenorhabditis elegans* feeds by drawing bacteria suspended in liquid into its pharynx, a neuromuscular organ that functions as a pump. The cycle of contraction and relaxation that draws food from the environment and filters bacteria from liquid is referred to as pharyngeal pumping. About one out of four pharyngeal pumps is followed by a posterior moving peristaltic contraction that transports food past the pharyngeal isthmus. This is referred to as isthmus peristalsis and, while coupled to pumping, it is distinctly regulated [2, 5, 9, 13]. The rate of pumping is thus the primary indicator of food intake [1].

The rate of pumping depends on feeding history, quality of food, and the familiarity of food [6, 12, 14]. Counting the stereotypical motion observed in the terminal bulb of the pharynx enabled detailed analyses of neuronal and molecular mechanisms that regulate pumping. However, these regulatory pathways were predominantly examined in a stationary environment, containing a saturated, high abundance of (familiar) bacteria on a standard agar plate. Moreover, traditional feeding assays rely on manual scoring of the mean number of pumps over brief (typically 30 sec) intervals [3, 10, 14]. The reduction of a potentially complex time-series to a single average rate may result in loss of pertinent information or even unintended bias.

Bacterial lawns on a standard cultivation plate are typically highly concentrated and large compared to the worm. Under these conditions, reported pumping rates are 4-5 Hz and the duration of a single pump (constrained by the physiology of the pharynx) is approximately 170 ms [1]. The pharynx is separated from the rest of the body by the basal lamina, an extracellular formation of connective tissue. It contains 20 muscle cells and 20 neurons. Of these, only the MC cholinergic motoneurons were found to be individually required for rapid pumping in the presence of food [3, 10]. The cholinergic neurons M2, M4, MC, and I1 form a degenerate network, excitatory for pumping and robust, where I1 can activate both MC and M2 [15]. In addition, the glutamatergic M3 neurons regulate the termination of a pump and M4 regulates isthmus peristalsis [1, 3, 10]. Under standard conditions, M3, M4, and MC are sufficient for supporting nearly normal feeding and growth [5, 10]. The functions of additional pharyngeal neurons are poorly understood. In part, this may be due to the challenges of characterizing the phenomenology of pharyngeal pumping more systematically.

Here we describe an affordable and scalable method for automatically assaying pharyngeal pumping. Our method combines a previously described microfluidic device [8], low cost educational microscopes, and an image analysis pipeline implemented using widely available open source tools and libraries [11]. The advantages of our approach include precise control of conditions such as the quality, uniformity, and concentration of available food, the possibility of prolonged measurement durations (hours, if required), unbiased automatic detection of pumping events, and the possibility to assay feeding conditions that change dynamically in a controlled manner. Manual scoring of pumping is arduous and limited to brief measurement periods. Automatic scoring of pumping on a high quality microscope is throughput-limited due to the cost of the imaging equipment. We found that lower quality imaging can be compensated for by rapid sampling and improved analysis without compromising the quality of the data. Using three microscopes we were able to assay up to 50 animals per day, i.e., 6 animals per objective, each for a full hour. The cost of the required equipment was $2, 000 (not including optional syringe pumps or other pressure sources). The setup can be further duplicated and the rate limiting step is the number of microfluidic devices that the researcher can load. The degree to which food availability can be dynamically controlled is dictated by the design of the microfluidic device used to restrict the motion of the animals and by the capabilities of the pressure source that is driving the fluid flow.

To validate our method we have compared the automatic detection to manual scoring. In addition, we characterized the phenomenology of feeding under three fixed food concentration, two velocities of fluid flow through the device, and in the presence of oscillations in food availability. We confirmed that the rate of pharyngeal pumping increases with the concentration of available food and that bursts of continuous pumping are interspersed with intervals of inactivity. In our hands, the rate of fluid flow affected the rate of pumping significantly but mildly. Finally, we measured pumping in the presence of oscillations in food availability. Interestingly, the mean rate of pumping was reduced two fold with respect to comparable steady state conditions. When pumping dynamics of individual animals were analyzed we found that, in addition to low mean rates, shifts in food concentration often failed to evoke correlated changes in the rate of pumping. Therefore, our data suggests that *C. elegans* feeding in a dynamic environment may exhibit correlations on timescales of several minutes.

## II. RESULTS

### A. A scalable imaging setup

We found that the image quality of commonly available educational microscopes, such as the Celestron 44104, is sufficient for assaying pharyngeal pumping. This model is equipped with 4x, 10x, and 40x air objectives, LED illumination, and a manual dual-axis translation stage. For convenience, a simple mount for the microfluidic device can be assembled from laser-cur Polymethyl methacrylate (PMMA, acrylic). Basler acA1920-25um scientific CMOS cameras were used for imaging. Mounting the cameras can be achieved by gluing a laser-cut acrylic ring to a C-mount adapter (Thorlabs SM1A9) or by 3D printing a suitable mount. Image acquisition was controlled by a small desktop computer equipped with a three drive RAID 0 array for fast storage. Animals were assayed on WormSpa microfluidic devices [8]. An Elveflow OB1 Mk3 pressure controller and and MUX-D10 automated injection valve were used to dynamically change food availability.

### B. Image analysis

Pharyngeal pumping motion is most easily detected through the motion of the grinder - a cuticle region of the pharynx inside the terminal bulb that is used to crush bacteria to aid digestion [4]. At low magnifications, directly tracking the position of the grinder is ineffective: the amplitude of grinder motion is small and any translation or deformation of the head will compromise the data. Therefore, our approach relied on intensity differences between consecutive images and on the separation of timescales between head motion and pumping (Fig. 1A-B). Key to this approach is a high imaging rate, 62.5 fps, more than 10 times faster than the maximal instantaneous rate of pumping, 6 Hz. The position of the head is maintained by mounting the animals into the WormSpa microfluidic device such that tracking a moving animal is not required and the motion of the head is dampened without a major impact on feeding [8].

**FIG. 1.**
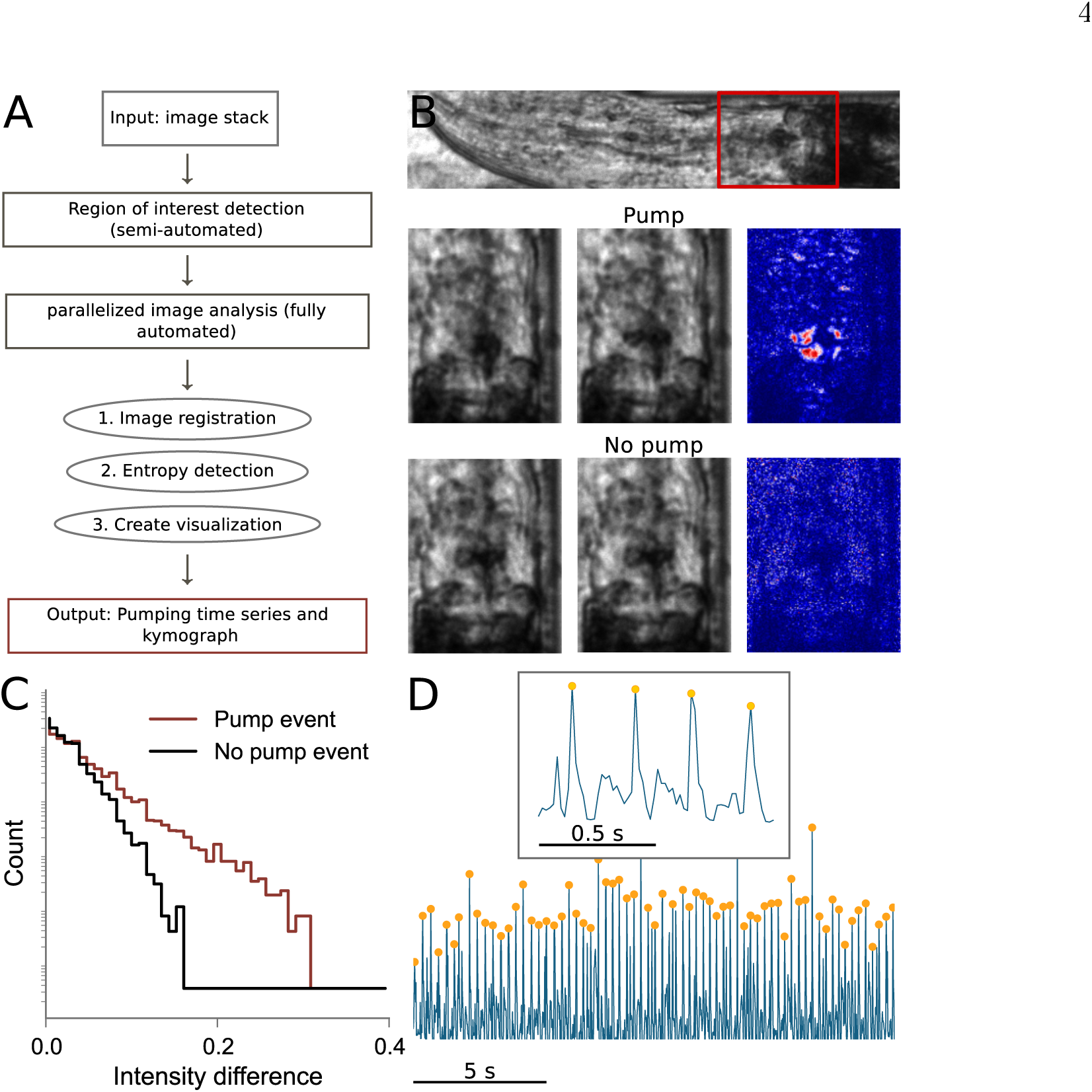
Entropy-based detection of pumping events. (A) An overview of the work flow from image acquisition to the time series of entropy peaks. Individual steps require little user input such that multiple animals can be analyzed in parallel. (B) Top: an image of the head of a nematode in a WormSpa microfluidic device mounted on an educational microscope (10x magnification). The terminal bulb is enclosed in a red box and the grinder appears as a dark spot at its center. Middle: a pair consecutive frames captured during a pump and a heat map of the intensities in the corresponding difference image. Bottom: same as above for a pair of frames captured in the absence of a pump. Pumping results in a small number of high intensity pixels in the difference image. (C) The distribution of intensity values in the difference image in the presence (red) and absence (black) of a pump. The separation between the two histograms is most pronounced at the high intensity tail of the distribution. (D) A sample trace showing the dynamics of entropy during four (top) or many (bottom) pumping events. individual events correspond to characteristic entropy peaks which are readily identified by standard peak detection algorithm followed by a minimal height selection criteria (orange dots) and the removal of close double peaks (less than ten frames apart).

Subtracting consecutive frames that were captured 16 ms apart isolates fast-changing features, i.e., the motion of the grinder. Examples of distributions of intensity differences in the presence or absence of a pump are shown in Fig. 1C. Notably, rapid motion affects the tail of this distribution. Therefore, a measure that preferentially weighs the tail would enhance the signal to noise ratio of motion detection. Using the entropy, Σ_*i*_ *p*_*i*_ log *p*_*i*_ (where *p*_*i*_ is the probability of observing intensity *i* in the difference image), to enhance the significance of motion achieves this goal [7]. An additional advantage of this method as compared to tracking the grinder is that no model of the background needs to be calculated. In a high frame-rate movie of grinder motion, the entropy of intensity differences peaks sharply when pumping occurs while contributions of slower head motions are minor. In our hands, imaging conditions did not require fine tuning in order to maintain a signal to noise ratio that exceeded 200% (1D-E).

Peaks in the entropy were identified using a standard peak detection algorithm. To validate the correspondence between such peaks and pumping events we compared the performance of our method to identifying pumps by visual inspection. The confinement of the animals constrains head motion to the direction of the longitudinal axis of the body. We thus generated kymographs of the animal over time by summing across the lateral axis of each frame. The grinder appeared as a dark spot in each line and pumps appear as stereotypical peaks. We verified that these peaks correspond to pumping motion in the raw images and counted kymograph peaks by visual inspection. We found that manual and automatic detection of pumping highly correlated (Fig. 2A). To compare our measurements to previously reported pumping rates, we binned our data in 30 sec intervals. The resulting pumping rates also exhibited good agreement between manual and automatic scoring (Fig. 2B). Likewise, the distributions of instantaneous rates, i.e., of the reciprocal values to the periods measured between consecutive pumps, were similar (Fig. 2C-D). Therefore, we concluded that our assays are capable of identifying pharyngeal pumping with a high level of accuracy.

**FIG. 2.**
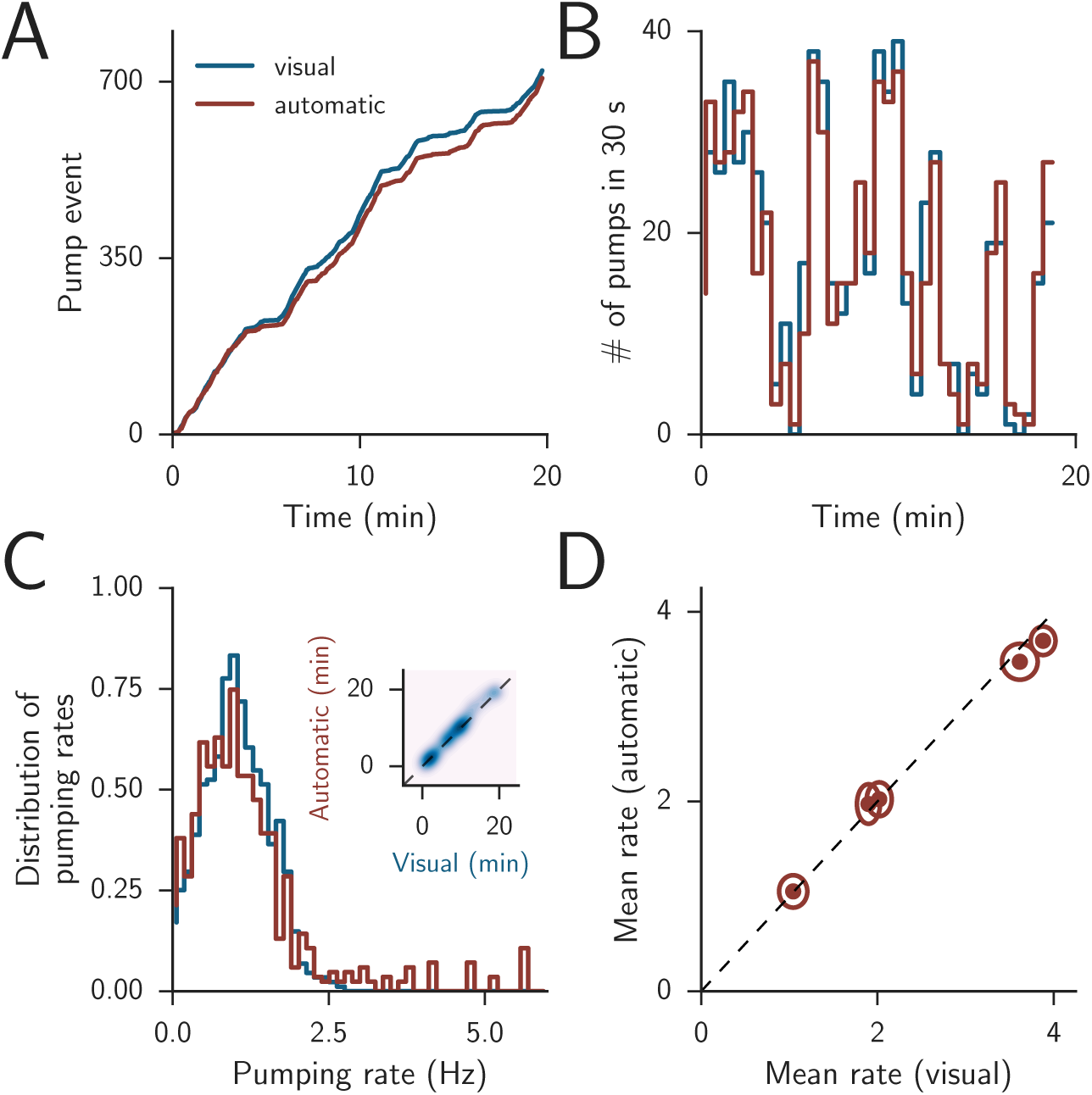
Pumping detection at low magnification (4x). All panels correspond to 20 min recordings of wild-type animals in which automated and manual scoring were compared. Unless stated otherwise, the concentration of bacterial food (*E. coli* OP50) was *OD*_600_ = 2.5. In panels A-C, automated and manual scoring are plotted as red and blue lines, respectively. (A) An example of the cumulative number of pumps as a function of time (*R*^2^ = 0.98). (B) The data from panel A binned in 30 sec bins (*R*^2^ = 0.90). (C) An example of a distribution of instantaneous pumping rates (intervals between consecutive pumps). Inset: correlation between the automatically detected and manually detected pump events. The dashed line denotes unity. Intensity corresponds to the density of pumping events. (D) Mean pumping rates from 5 animals (one per data point). Here, the instantaneous pumping rates were smoothed with a median filter of width 15. The dashed line denotes unity, the major and minor axes of the ellipsoids represent the standard deviations of the two rates. To obtain a range of mean rates, we assayed 5 animals at food concentrations between *OD*_600_ = 3 and *OD*_600_ = 5 and scored 500 pumping events per animal.

### C. Pharyngeal pumping is affected by the concentration and the flow rate of bacterial food.

To demonstrate our method, we first assayed pharyngeal pumping in the presence of fixed food concentrations. In agreement with previous reports, pumping rates decreased when the concentration of bacteria was lowered but did not vanish in the absence of food (Fig. 3A-C). In addition, the mean rates we observed were comparable (Fig. 3C) to the values reported previously [1, 6, 10]. Henceforth, unless stated otherwise the high and low food concentrations were set to *OD*_600_ = 0.0 and 4.0, respectively. Fig. 3A-B depict typical distributions of instantaneous pumping rates under these conditions. When food was readily available, wild-type animals exhibited a peak rate of 4 — 5 Hz but also a substantial tail of lower rates. In our hands, changing the flow rate in the device from 5 to 200 *µl/min* resulted in a mild yet significant reduction in pumping rates (Fig. 3A, C). Our measurements were performed continuously and therefore invariably included periods devoid of pumping events, as indicated by the low frequency tail in Fig. 3A. Therefore, we also measured the duration of continuous pumping, defined as a series of consecutive pumps that were separated by no longer than 250 ms. The fraction of time spent in continuous pumping was defined as the pharyngeal duty ratio, which also increased with food concentration (Fig. 3D). These results indicate that our method provides an unbiased and efficient approach for characterizing pharyngeal pumping at precisely controlled steady state conditions.

**FIG. 3.**
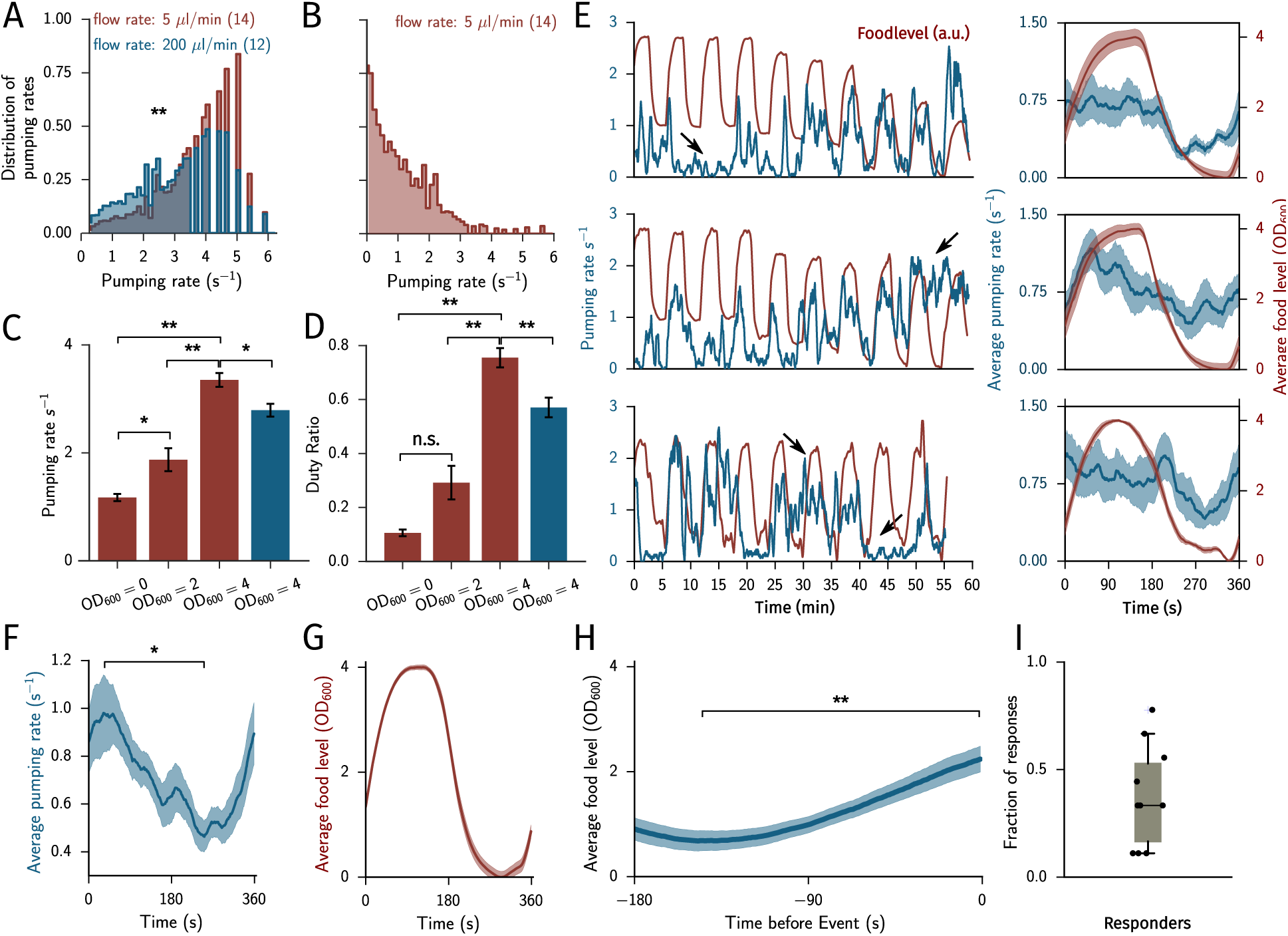
Pumping at variable food levels. All panels correspond to recordings from wild-type animals at 4x magnification. In panels A-D each animal was assayed for 30 min in the presence of a constant concentration of food (*E. coli* OP50). In panels E-I each animal was assayed for 60 min. Pumping rates and food levels are denoted in blue and red, respectively. Bacterial food flowed at a rate of 200 *µl/min* and oscillated between high (*OD*_600_ = 4.0) and low (*OD*_600_ = 0.0) levels with a period of 360 sec. (A) Distributions of wild-type instantaneous pumping rates at a food concentration of (*OD*_600_ = 4.0 were affected by the flow rate in the device. Flow rates of 5 *µl/min* and 200 *µl/min* are plotted in red and blue, respectively. *N*_5 *µl/min*_ = 14, *N*_200 *µl/min*_ = 12 animals (B) The distribution of instantaneous pumping rates in the absence of bacterial food (*OD*_600_ = 0.0). *N* = 14 (C-D) Mean pumping rates and duty ratios for the data presented in panels A-B. The duty ratio was defined as the fraction of time occupied by continuous pumping, i.e., when consecutive pumps were ≤ 250 ms apart. *N*_*OD*=0_ = 14, *N*_*OD*=2_ = 12, *N*_*OD*=4, 5*µl/min*_ = 14, *N*_*OD*=4_, _200*µl/min*_ = 12. (E) Left: representative examples of pumping dynamics of individual animals in the presence of oscillating food availability. Food concentration is denoted in red. Pumps were grouped in bins of 1 *sec* and the time-series was smoothed using a moving average with 20 *sec* width. Arrows denote periods where the animal maintained high or low pumping activity regardless of the oscillations in food concentration. Right: the mean pumping rates and food levels per animal, i.e., data were averaged over the corresponding nine stimuli for each of the animals on the left. (F-G) Mean pumping rates and food levels averaged over stimuli and all animals. Shaded areas denote ± s.e.m. (H) Response-triggered average to the periodic stimuli: food concentration was averaged during all 180 sec periods that preceded time-points where the pumping rate exceeded 2 Hz (defined as *t* = 0). The blue line depicts the population mean and the shaded area denote ± s.e.m. (I) Fraction of responses per animal. A response to a stimulus was defined as an increase in pumping rate during high food levels that was at least 2 s.e.m higher than the mean rate at the preceding *OD*_600_ = 0.0 period. In panels F-I *N* = 10 animals.

### D. Pumping rates are reduced and exhibit long correlations in a dynamic environment.

Combined, controlling food availability and prolonged measurements enable assaying pharyngeal pumping in a dynamic environment. To demonstrate this, we have oscillated the concentration of available food between its high (*OD*_600_ = 4.0) and low (0.0) values with a period of 360 sec. Switching between the two food reservoirs was performed using a small volume rotative valve, which eliminates back-flow and cross contamination and minimizes flow disruptions. As a result, we could not detect a response to switching between two reservoirs containing the same concentration of food (data not shown).

Panels E-G in Fig. 3 depict examples of instantaneous pumping rates of individual animals and the corresponding food levels, as well as averaged data. Our analysis could readily detect changes in pumping rates that occurred over ≤ 10 sec and in this assay the animals experienced a plateau of high food availability for 90 sec during each period. We thus hypothesized that pumping rates would track the changes in food availability and that their highest and lowest mean pumping rates would be comparable to those measured at the corresponding fixed concentrations (Fig. 3C). However, mean pumping rates in the dynamic environment were approximately 2-fold lower than the corresponding steady state rates (Fig. 3F-G). Moreover, it was not uncommon for animals to maintain low or high pumping activity regardless of oscillations in the concentration of food for up to 10 min (see arrows in Fig. 3E and Fig. 3I).

Due to the variability of the behavioral responses, we also calculated the response-triggered average to the periodic stimuli. We identified the time points when the instantaneous pumping rate exceeded a threshold value of 2 Hz. Food concentration data from the 180 sec periods that immediately preceded these time points were aligned and averaged. We found that the mean concentration of food increased from *OD*_600_ ⋍ 0.7 at earlier times to *OD*_200_ ⋍ 2.2 when high activity was detected (Fig. 3H). This result indicates that high pumping rates are exercised preferentially in response to an increase in food availability. Combined, our results suggest that *C. elegans* readily detects dynamic changes in food availability and can respond to them rapidly, yet their behavior is more complex than simply tracking the external conditions.

### III. DISCUSSION

We described an efficient, unbiased, and scalable method for measuring pharyngeal pumping of *C. elegans* under controled and reproducible conditions. Our method exhibits good agreement with previously reported data under comparable conditions, as well as good agreement between manual and automatic scoring of pumps. Moreover, it extends the conditions of pumping assays to prolonged measurements and dynamic yet controlled environments. Consequently, we found that *C. elegans* responds to dynamic changes in food availability with unexpected complexity: the overall rate of pumping is suppressed and periods of high or low pumping activity can persist for several minutes regardless of the changes in available food.

As compared to locomotion-based behaviors, extensively studied under a wide range of circumstances, pharyngeal pumping assays were by and large performed under a restricted set of conditions. The combination of rapid and spatially minuscule motion limited the throughput of pumping assays, shortened the duration of individual measurements, and served as an additional barrier to challenging the animal with dynamic environments. Therefore, our approach can be used to address a variety of questions that were minimally studied previously. These include the quantitative relations between food availability and the instantaneous rate of pumping or the impact of external dynamics on pumping responses.

Our methodology can be easily adapted and modified to accommodate varying experimental needs. For instance, the image analysis can be combined with a worm tracker (at the expense of throughput) to assay pumping in freely locomoting animals and alterations to the microfluidic devices can greatly enhance the flexibility of the assay. At its heart, our approach trades image quality for frame rate while preserving the integrity of the data. Due to its low cost, it can be considerably scaled up in size. As the cost of electronics such as low-end desktop computers and CMOS cameras continues to drop, the limiting factor would likely be the schedule of the researcher performing the assays.

## IV. ADDITIONAL MATERIALS AND METHODS

### A. Animal handling

*C. elegans* N2 animals were cultivated with OP50 bacteria according to standard protocols at 20°C. To measure the statistics of pumping in detail, we use a microfluidic device that enables continuous measurements of pumping and control of the feeding conditions [8]. Young adults were picked into liquid NGM and loaded into the WormSpa microfluidic device. An *E. coli* OP50 overnight culture, concentration-adjusted to the desired *OD*_600_ in NGM was flown through the device at a constant rate throughout the assay. After an hour of acclimation in the device, the animals were imaged for one hour at a magnification of 4x and 62.5 frames per second using the optical setup.

### B. Monitoring oscillating food levels

Food levels were oscillated using a programmable pressure regulator (Elveflow Inc., Paris, France). High and low food levels were measured prior to the assay. Food coloring ( McCormick & Co., Hunt Valley, 21031, MD) was filtered through a 0.22 *µ*m filter and diluted to a final concentration of 1: 60 in the tube containing the low food concentration. In preliminary assays we observed that oscillating between two sources containing identical concentrations of food, one of which supplemented with food coloring, did not evoke a correlated response (data not shown). Thus, our assay could not detect a response to neither the food coloring nor the switching between the two sources. The food coloring shifted the mean brightness of our images, which indicated the concentration of food flowing through the device, without obscuring the anatomy of the pharynx.

### C. Statistics

In Figs 3C, D, F and H significance was calculated using an unequal-variance t-test and the Bonferroni post-hoc correction for multiple comparisons (3C and 3D). In 3F and 3G values at the minimum and maximum of the average curves were compared. The distributions in 3A were compared using a *χ*^2^-test with 35 classes (non-zero bins).

